# Mapping the sex determination region in the *Salix* F_1_ hybrid common parent population confirms a ZW system in six diverse species

**DOI:** 10.1101/2021.11.04.467334

**Authors:** Dustin G. Wilkerson, Bircan Taskiran, Craig H. Carlson, Lawrence B. Smart

## Abstract

Within the genus *Salix*, there are approximately 350 species native primarily to the northern hemisphere and adapted to a wide range of habitats. This diversity can be exploited to mine novel alleles conferring variation important for production as a bioenergy crop, but also to identify evolutionarily important genes, such as those involved in sex determination. To leverage this diversity, we created a mapping population by crossing six *Salix* species (*S. viminalis, S. suchowensis, S. integra, S. koriyanagi, S. udensis*, and *S. alberti*) to common male and female *S. purpurea* parents. Each family was genotyped via genotyping-by-sequencing and assessed for kinship and population structure as well as the construction of 16 backcross linkage maps to be used as a genetic resource for breeding and selection. Analyses of population structure resolved both the parents and F_1_ progeny to their respective phylogenetic section and indicated that the *S. alberti* parent was misidentified and was most likely *S. suchowensis*. Sex determining regions were identified on *Salix* chromosome 15 in the female-informative maps for seven of the eight families indicating that these species share a common female heterogametic ZW sex system. The eighth family, *S. integra* × *S. purpurea*, was entirely female and had a truncated chromosome 15. Beyond sex determination, the *Salix* F_1_ hybrid common parent population (*Salix F*_*1*_ HCP) introduced here will be useful in characterizing genetic factors underlying complex traits, aid in marker-assisted selection, and support genome assemblies for this promising bioenergy crop.

## INTRODUCTION

The establishment of genomic resources is an important step in developing a fully realized breeding program, reinforced by modern tools for trait mapping, candidate gene identification, and marker-assisted selection. *Salix* and *Populus* (poplars) comprise the majority of species in the family Salicaceae, which consists of dioecious trees, shrubs, and subshrubs that are highly heterozygous. Shrub willow (*Salix* spp.) are grown in northern latitudes as a sustainable, high-yielding, carbon neutral, bioenergy crop that can grow on marginal land and provide multiple ecosystem services (Smart et al. 2005; Stoof et al. 2014; Clifton-Brown et al. 2019; Fabio and Smart 2020). While shrub willow breeding in the United States has been active since the 1980’s, most of the nearly 350 species have yet to be tapped as a source of genetic diversity (Dickmann and Kuzovkina 2014; Stanton et al. 2014).

Genomic resources developed for poplar were used in early genomic studies in *Salix*, because their genomes are largely co-linear (Hanley et al. 2006; Berlin et al. 2010). As sequencing technologies became increasingly more affordable, more *Salix* genome sequencing was completed and now there are several high-quality assemblies. Genomic resources for *Salix* are currently centered around a few key species. In Europe, *Salix viminalis* is an important bioenergy crop species with a recently published, high quality genome assembly (Almeida et al. 2020). *S. viminalis* has been used in several QTL mapping studies for resistance to willow leaf rust (*Melampsora larici-epitea*) (Rönnberg-Wästljung et al. 2008; Samils et al. 2011; Sulima et al. 2017), drought tolerance (Rönnberg-Wästljung et al. 2005), and growth and phenology (Hallingbäck et al. 2016; Hallingbäck et al. 2019). While in the United States, *Salix purpurea* is the model species for bioenergy willow breeding, genetics, and genomics. The US Department of Energy Joint Genome Institute has produced the highest quality annotated *Salix* reference genomes assembled in the genus on male and female *S. purpurea*, available on Phytozome (Zhou et al. 2018; Zhou et al. 2020) (https://phytozome-next.jgi.doe.gov/info/SpurpureaFishCreek_v3_1 https://phytozome-next.jgi.doe.gov/info/Spurpurea_v5_1). Using joint linkage and association mapping approaches focused on *S. purpurea*, Carlson et al. (2019) identified numerous QTL for a wide range of morphological, physiological, insect and disease resistance and biomass composition traits. A naturalized species in North America, *S. purpurea* is a potential donor of broad adaptability traits for species susceptible to pests and diseases in the Northeast United States. There is value in studying the genomes of less-characterized *Salix* species for phylogenomic analysis and to discover diverse sources of alleles for introgression into elite yielding cultivars.

Here, we introduce the *Salix* F_1_ hybrid common parent population (*Salix* F_1_ HCP). The parents, described in Fabio et al. (2019) and Crowell et al. (2020), represent a diverse selection of species from *Salix* subgenus Vetrix (Dickmann and Kuzovkina 2014). Six *Salix* species (*S. viminalis, S. suchowensis, S. integra, S. koriyanagi, S. udensis*, and *S. alberti*) were crossed to common male and female *S. purpurea* parents to form eight species hybrid families. Literature describing these species ranges from the high-quality reference genomes available for *S. purpurea* and *S. viminalis* to the scarcely studied *S. alberti* (Rosso et al. 2013). *Salix suchowensis* is native to China and has been used recently to generate a chromosome scale genome assembly (Wei et al. 2020). This species has been assessed for its response to drought stress (Jia et al. 2020) and was one of the first *Salix* species used to map the sex determination region (SDR) (Liu et al. 2013). *Salix udensis*, formally known as *S. sachalinensis*, has been described as a Japanese riparian willow species that acts as a natural nest cavity for fish owls (Niiyama 2008; Slaght et al. 2018) and is suggested to have sexually dimorphic characteristics (Ueno and Seiwa 2003; Ueno et al. 2006). Some genomic resources are available for the Korean *S. koriyanagi*, as its chloroplast genome have been sequenced (Kim et al. 2019; Park et al. 2019) while *S. integra* has been assessed for its phytoremediation potential (Cao et al. 2020; Yang et al. 2020b). Developed to interrogate the genetics of several understudied *Salix* species, the *Salix* F_1_ HCP is an important step in the development of genomic resources in *Salix*.

The Salicaceae represents an interesting family for the study of the evolution of dioecy and the mechanisms of sex determination. Genetic mapping of SDRs in the Salicaceae have revealed considerable intra- and inter-chromosomal variability and contrasting sex determination systems (Yang et al. 2021). While *P. alba* and *P. trichocarpa* are male heterogametic (XY) with SDRs located on chr19, the SDR of *P. euphratica* was identified on chr14 and is female heterogametic (ZW) (Paolucci et al. 2010; Geraldes et al. 2015; Yang et al. 2021). Conversely, the SDR of *S. purpurea* has been mapped to a large, pericentromeric region on *Salix* chr15, but has been found to maintain a few orthologous regions present within the *P. trichocarpa* chr19 SDR (Zhou et al. 2018). However, it is unclear whether dioecy evolved before or after their divergence (Hou et al. 2015). The sex systems of *Salix* also lack a consensus model. For instance, similar to *S. purpurea*, both *S. suchowensis* (Hou et al. 2015; Chen et al. 2016) and *S. viminalis* (Pucholt et al. 2015; Pucholt et al. 2017b) have female heterogametic (ZW) systems with SDRs located on *Salix* chr15, yet tree-form *S. nigra* has an XY system SDR on chr07 (Sanderson et al. 2021). Considering the variability in sex determination systems and SDR locations already discovered within *Salix*, elucidating the mechanisms of sex determination in more species could help to build a more cohesive understanding of SDR evolution. Further investigation could provide evidence to implicate SDR turnovers as a factor in the considerable species diversity within the genus.

To establish the *Salix* F_1_ HCP as a genetic resource, this study sought to: (1) describe the genetic relationships between the parent species, (2) develop female and male parent informative linkage maps for each family, and (3) define and compare their respective SDRs.

## MATERIALS AND METHODS

### Population Development

Females of *S. viminalis, S. integra, S. alberti*, and *S. suchowensis* were crossed with male *S. purpurea* clone ID 94001, while males of *S. viminalis, S. udensis, S. koriyanagi*, and *S. suchowensis* were crossed with female *S. purpurea* clone ID 94006 creating eight F_1_ species hybrid families (Fig. 1). The common *S. purpurea* parents, 94006 and 94001, were chosen based upon their differential resistance to willow leaf rust (Crowell et al. 2020), adaptation to the Northeastern U.S., and availability of high-quality reference genomes (Zhou et al. 2020).

**Figure 1:**
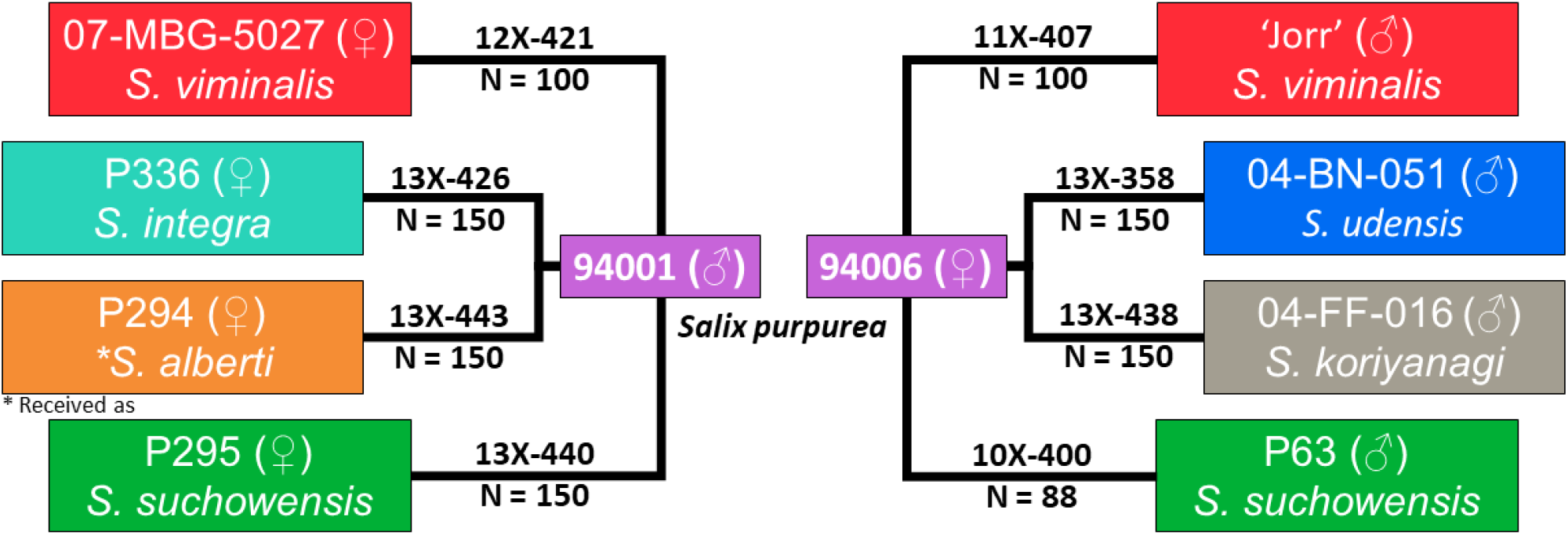
Pedigrees of the *Salix* F_1_ HCP. There were four full-sib families with 94001 as the paternal parent and four full-sib families with 94006 as the maternal parent. Reciprocal crosses were made with male and female *S. viminalis* and *S. suchowensis* while *S. integra, S. alberti, S. udensis*, and *S. koriyanagi* were crossed only once. P294 was received with the identification of *S. alberti*.

Crosses were made by forcing floral catkins of the parent genotypes from dormant shoots in a greenhouse. All crosses were made in isolation to prevent pollen contamination. When anthers began to dehisce, male catkins were excised and placed in falcon tubes for pollen extraction using toluene, as described in Kopp et al. (2002), then stored in 2 mL microcentrifuge tubes at -20°C until female catkins were receptive. Seedlings were established in a standard peat-based potting mix in a greenhouse, then transplanted to nursery beds near Cornell AgriTech (Geneva, NY). Winter dormant cuttings from all parents and progeny were collected from one-year old stems and hand-planted in the field in a randomized complete block design with three plants per plot and four replicate blocks. Field trials were established with 1.83 m spacing between rows and 40.6 cm between plants within rows.

### DNA Extraction and Genotyping-by-Sequencing

Shoot tips for DNA extraction were collected from plants in nursery beds and stored in desiccant. Dried shoot tips were ground to a fine powder with a Geno/Grinder (SPEX SamplePrep, Metuchen, NJ, USA) prior to genomic DNA extraction using the DNeasy Plant Mini Kit (QIAGEN Inc., Valencia, CA, USA). After checking the DNA quality using gel electrophoresis, DNA quantity was estimated using the Invitrogen Qubit dsDNA Broad Range Assay kit on a Qubit Fluorometer (ThermoFisher Scientific, Waltham, MA). Genomic DNA was submitted to the University of Wisconsin Biotechnology Center (Madison, WI) for 96-plex Genotyping-by-Sequencing (GBS) library preparation using the *ApeK*I restriction enzyme and sequenced (1 × 100 bp) on the Illumina HiSeq 2500 (Illumina, Inc., San Diego, CA, USA) platform.

### Variant Discovery and Imputation

Initial variant discovery and filtering was performed using the TASSEL GBS v2 Discovery Pipeline (Bradbury et al. 2007). Sequence reads were trimmed to 64 bp and aligned using BWA mem (Li and Durbin 2009) under default parameters to the *S. purpurea* v5.1 reference genome (Zhou et al. 2020). As this genome includes both chr15Z and 15W, chr15Z was excluded to reduce mapping errors within the pseudoautosomal regions surrounding the SDR in *S. purpurea*. This process was repeated once for all eight families together and then for each of the eight F_1_ families separately to identify population-wide and family-specific SNPs. By calling variants on all eight families and then on each family separately, variants were based on inter- and intra-familial genetic differences, respectively. The resulting VCF files contained 684,412 SNPs for the full analysis and ranged from 174,762 to 266,797 SNPs depending on the family. On the eight F_1_, SNPs with > 70% missing data and minor allele frequency < 0.01 and F_1_ individuals were dropped if they had > 80% missing data or determined to be outliers based on principal component analysis. Missing genotype calls and low read depth are common in GBS (Elshire et al. 2011), therefore imputation was performed separately on each family to validate calls in haplotype blocks. Using LinkImputeR (Money et al. 2017), genotypes called with a read depth < 5 were set to missing before filtering again for missingness > 70%, resulting in 95,281 to 145,944 imputed SNPs with accuracies ranging from 84.3 to 93.1%.

### Population Structure

SNPs were filtered to retain markers and individuals with ≤ 20% missing data and SNPs with a minor allele frequency > 0.01, which resulted in 55,398 SNPs. Analysis of principal components and kinship using default parameters were performed in Tassel 5 and visualized in R (R Core Team 2020). Multiple runs of the parents and 10 randomly selected F_1_ progeny were analyzed using fastSTRUCTURE (Raj et al. 2014). Only a subset of the F_1_ were used in this analysis in order to manage file size and computation requirements. Multiple analyses of ‘structure.py’ were completed (K = 3 - 10). Using ‘chooseK.py’, K = 6 represented the model complexity that maximized the marginal likelihood and best explained the data structure, suggesting six separate populations.

### Linkage Map Construction and Analysis

Unless otherwise indicated, linkage map construction and analysis were performed using custom R code, available on Github (link in Data Availability). Multiple runs of each parent were used to form consensus genotypes for each SNP. In the absence of a clear consensus, the genotype was set to missing. If both parents were set to missing, the SNP was removed from the analysis. If only one parent was set to missing, its genotype was inferred based on the genotype of the known parent and the segregation of the F_1_. These parental consensus genotypes were used to identify the female informative (AB × AA) and male informative (AA × AB) markers used to generate backcross linkage maps for each parent using a combination of R/qtl (Broman et al. 2003) and ASMap (Taylor and Butler 2017). Co-located markers and those exhibiting extreme segregation distortion were removed using ASMap function ‘pullCross’.

Linkage groups were then created using ‘mstmap’ with default parameters except for objective.fun = “ML” and bychr = FALSE. The p-value for determining linkage groups varied between families, ranging from 1e-6 to 1e-12, depending on the demarcation of the 19 expected linkage groups. A custom R function was then used to perform simple error correction to reduce the number of double crossovers and deflate map distances before reforming linkage groups. Briefly, this function relies on the marker order determined after formation of linkage groups, identifying double crossovers of ≤ two SNPs, and correcting them. Map quality was checked using two strategies. The ‘heatMap’ function in ASMap, which plots LOD linkage between markers on the upper triangle and estimated recombination frequency on the bottom, reveals markers that are problematic or out of phase. Then by comparing the physical position (Mb) based on alignment to the *S. purpurea* reference genome and the genetic distance (cM) within each linkage group shows issues with marker order or potential chromosomal rearrangements.

To delimit the SDR in each family, the sex of all F_1_ individuals was recorded by inspecting flowering catkins over two growing seasons in the field. Sex ratio bias was tested using a chi-square test for a 1:1 sex ratio. Using functions in R/qtl, genotype probabilities were calculated using ‘calc.genoprob’ (step = 0, error.prob = 0.01, map.function = “kosambi”, stepwidth = “fixed”). Next, QTL mapping for individual sex was performed using ‘scanone’: model = “binary”, method = “em”. Genome-wide significance thresholds were determined based on the results of ‘scanone.perm’ (n.perm = 1000). QTL positions were refined using ‘refineqtl’ (method = “hk”, model = “binary”), then the 1.5 LOD support intervals were calculated using ‘lodint’ with default parameters.

## RESULTS

### Population Structure

A combination of PCA, hierarchical clustering, and fastSTRUCTURE were used to describe the population structure of the *Salix* F_1_ HCP. A PCA of the parents revealed three distinct clusters, formed by two PCs accounting for 36.8% and 22.5% of the total genetic variation (Fig. 2A). The two common *S. purpurea* parents, 94006 and 94001, formed a single group and were separated from the other parents by PC2. The *S. viminalis* parents, ‘Jorr’ and 07-MBG-5027, formed a cluster with *S. udensis* 04-BN-051, which was differentiated from the remaining species by PC1. Including the F_1_ progeny from each family into the analysis, each PC accounted for 26% and 10.1% of the total variation (Fig. 2B). PC2 split *S. udensis* from the two *S. viminalis* parents and separated *S. koriyanagi* from *S. suchowensis, S. integra*, and *S. alberti*. As expected, the F_1_ individuals were intermediate between the common parent and family specific species. The F_1_ progeny derived from the female parents *S. suchowensis* P295, *S. alberti* P294, and *S. integra* P336 (all crossed with male *S. purpurea* 94001), co-localized.

**Figure 2:**
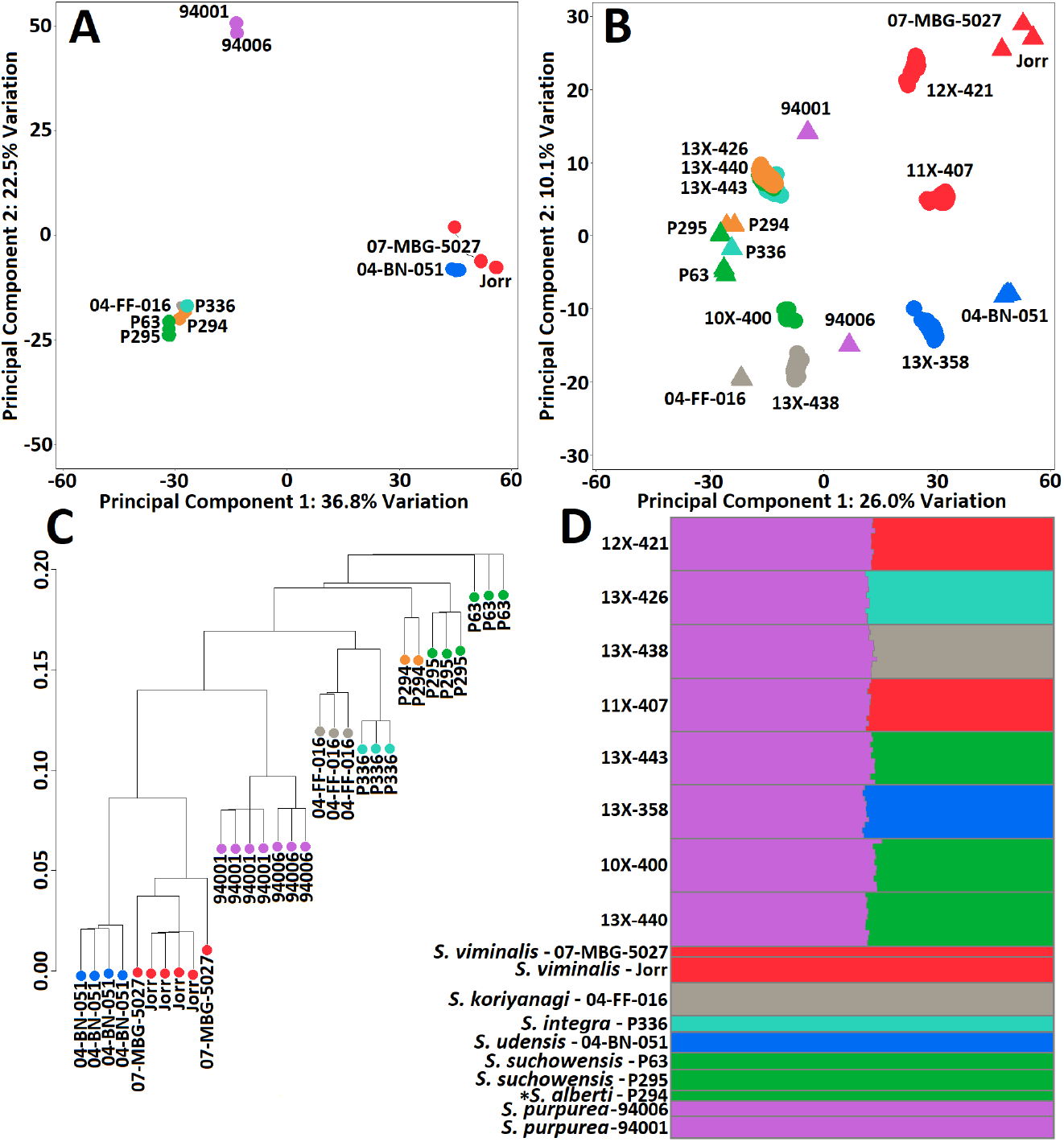
Results of PC and fastSTRUCTURE analysis of the *Salix* F_1_ HCP. A: PCA of the parents; B: PCA of the F_1_ and the parents; C: Hierarchical clustering of the parents; D: Distruct plot using fastSTRUCTURE results. In panels A, B, and C: purple - *S. purpurea*, red - *S. viminalis*, green - *S. suchowensis*, blue - *S. udensis*, teal – *S. integra*, grey – *S. koriyanagi*, orange - *S. alberti*. Colors in panel D are based on fastSRUCTURE results, affecting only *S. alberti*. *Received as *S. alberti*.

Kinship analysis grouped the two *S. viminalis* parents with *S. udensis*, the two *S. purpurea* parents together, and *S. koriyanagi* with *S. integra* (Fig. 2C). *Salix suchowensis* and the *S. viminalis* and *S. udensis* group were the least similar while *S. purpurea* grouped centrally between them, similar to PC1 (Fig. 2A). Within *S. suchowensis*, the female P295 was more closely related to the female *S. alberti* P294 than it was to P63, a male. Admixture analysis included the parents and a subset of the F_1_ individuals of each family. The eight groups of F_1_ individuals were comprised of roughly half the genetic background of *S. purpurea* and half the other species parent, as expected (Fig. 2D). *Salix viminalis, S. integra, S. koriyanagi, S. udensis* and *S. purpurea* formed distinct clusters, while *S. alberti* P294 grouped together with *S. suchowensis*.

### Linkage Map Construction and Analysis

Since there is considerable divergence between parents and pedigrees, variant discovery and marker filtration were performed for each family separately. Using consensus genotypes derived from multiple sequencing runs of the parents, markers were split into female (AB × AA) and male (AA × AB) informative backcross markers for linkage map construction, which resulted in 16 linkage maps (Fig. 3). Each linkage map consisted of 19 linkage groups with total map lengths ranging from 3939.9 – 6957.3 cM containing between 2035 and 3852 total markers (Table S1). Recombination frequency and genetic to physical distance plots generated for each linkage map revealed that marker order and phase within each linkage group were reasonably linear (Fig. S1). Sex phenotypes were used for QTL mapping of the SDR. Six of the eight families displayed significant sex ratio bias towards females based on a simple chi-square test (p < 0.05), with female to male ratios ranging from 1.4 to 1.6 (Table 1). Neither of the *S. viminalis* families displayed sex ratio bias, while the *S. integra* × *S. purpurea* family was entirely female. Single QTL for sex were identified on chr15 within seven of the eight maternal maps, excluding *S. integra* of 13X-426, explaining between 60.7 – 74.1% of the total phenotypic variation (Table 2). LOD scores for the female and male linkage maps are in Table S2 and Table S3, respectively. Genome-wide LOD significance thresholds ranged from 3.39 – 3.69, depending on the map. On the four maternal *S. purpurea* maps, the QTL as determined by the permuted significance thresholds accounted for 10.3 – 12.03 Mb of the roughly 15.5 Mb chr15 due to suppressed recombination (Figure 4). QTL refinement and 1.5 LOD support intervals narrowed this down to account for 1.17 – 7.31 Mb with peak markers located at 2.91, 2.92, 7.33, and 8.65 Mb when *S. suchowensis, S. viminalis, S. koriyanagi*, and *S. udensis* were the paternal parent, respectively (Table 2; Figure 4). Of the three remaining maternal maps, QTL determined by LOD significance accounted for 12.3, 11.45, and 13.55 Mb of chr15 on the *S. alberti, S. suchowensis*, and *S. viminalis* maps, respectively, reflecting suppressed recombination. Upon refinement, these ranges were narrowed to account for 1.59, 6.91, and 6.41 Mb of the chromosome, with peaks centered at 9.73, 2.57, and 2.92 Mb.

**Table 1:**
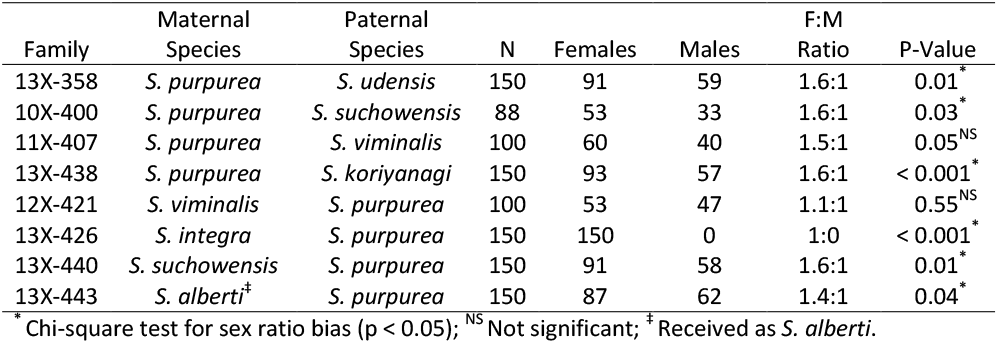
Sex phenotype statistics for the eight families in the *Salix* F_1_ HCP.

**Table 2:** Sex QTL associated with the SDR within the maternal linkage maps.

| Family | Maternal Species | Chr | Peak (cM) | Peak LOD | Peak (Mb) | Low-LSI (Mb) | High-LSI (Mb) | PVE (%) |
| --- | --- | --- | --- | --- | --- | --- | --- | --- |
| 13X-358 | *S. purpurea* | 15 | 85.3 | 29.3 | 8.65 | 4.51 | 9.22 | 73.7 |
| 10X-400 | *S. purpurea* | 15 | 81.1 | 16.1 | 2.91 | 2.61 | 9.92 | 61.8 |
| 11X-407 | *S. purpurea* | 15 | 114.2 | 18.6 | 2.92 | 2.88 | 4.05 | 60.7 |
| 13X-438 | *S. purpurea* | 15 | 70.1 | 35.1 | 7.33 | 2.63 | 8.29 | 71.1 |
| 12X-421 | *S. viminalis* | 15 | 86.0 | 21.1 | 2.92 | 2.95 | 9.36 | 66.0 |
| 13X-440 | *S. suchowensis* | 15 | 78.4 | 26.7 | 2.57 | 2.44 | 9.35 | 74.1 |
| 13X-443 | *S. alberti*<sup>‡</sup> | 15 | 145.8 | 35.6 | 9.73 | 8.63 | 10.22 | 70.3 |

LSI: 1.5 LOD Support Interval; PVE: Percent Variation Explained; cM: centiMorgan position within the linkage map; Mb: Physical position based on alignment to *S. purpurea* reference genome. <sup>‡</sup> Received as *S. alberti*.

**Figure 3:**
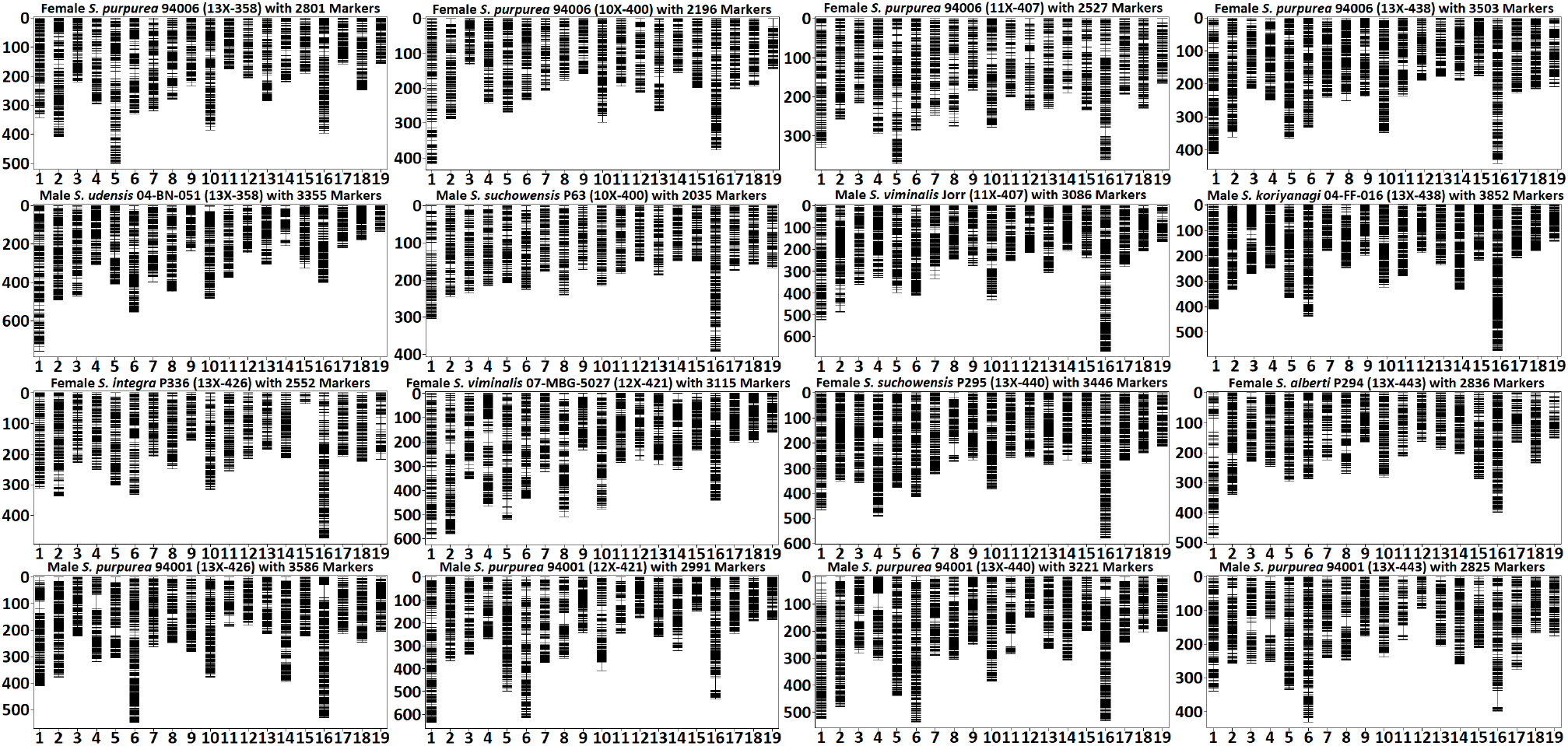
Linkage maps for each of the parents within the *Salix* F_1_ HCP. Female maps (first and third rows) were constructed using female informative markers (AB × AA), while male maps (second and forth rows) were constructed using male informative markers (AA × AB).

**Figure 4:**
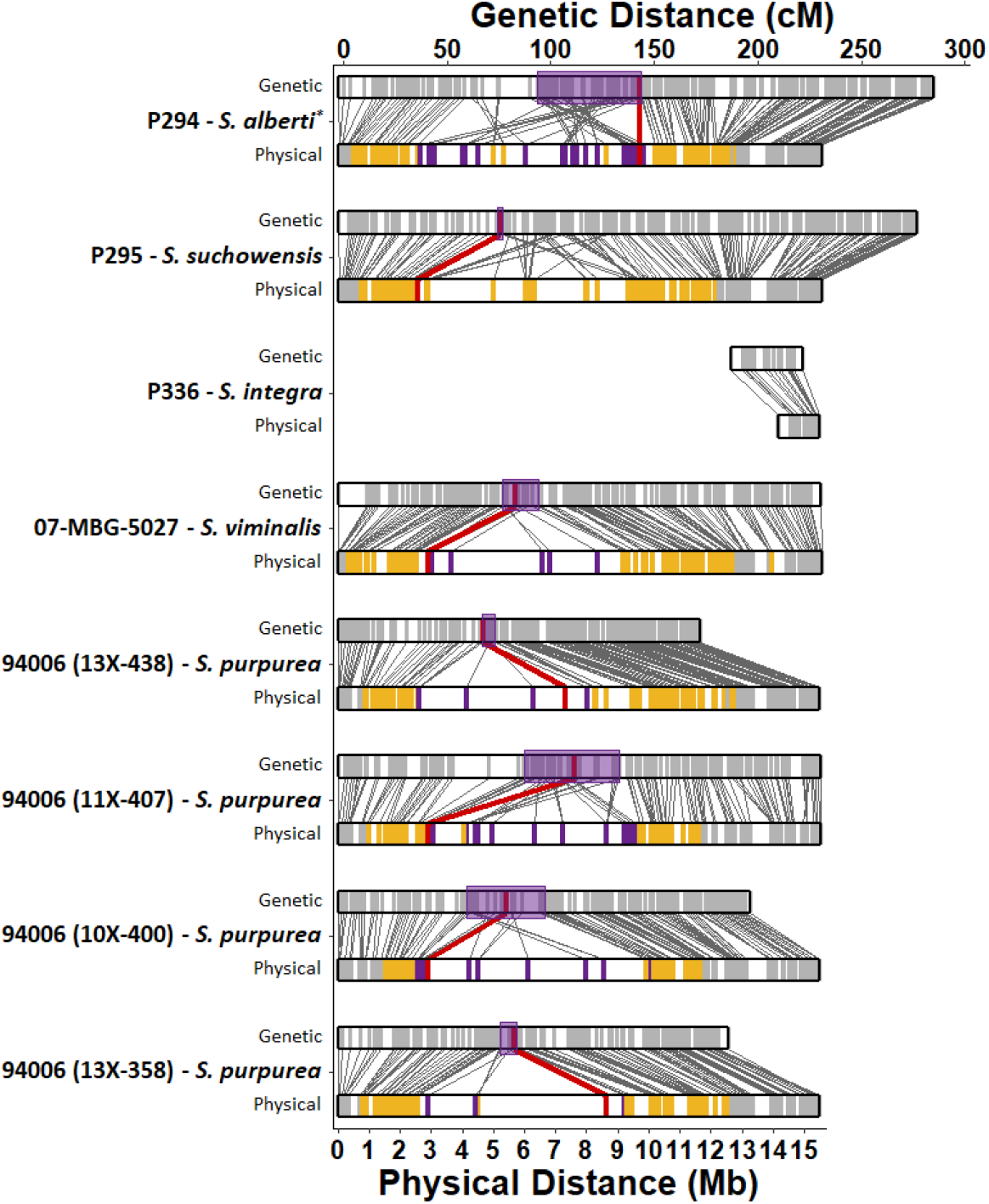
Genetic (cM) and Physical (Mb) distances for the maternal parents in the *Salix* F_1_ HCP. Each parent has a set of two maps, genetic on the top and physical on the bottom with lines connecting them indicating a marker’s relative position in both maps. On the genetic maps, each marker is represented by a grey vertical line while the 1.5 LOD support interval of the QTL is shown with a purple box. On the physical map, grey vertical lines are markers not associated with the QTL, yellow shows markers whose LOD score was above the permuted significance threshold, purple are markers within the 1.5 LOD support interval, and red indicates the cM and Mb position of the peak marker. P336’s linkage map cM distance was artificially increased for figure clarity.* Received as *S. alberti*.

## DISCUSSION

*Salix* is a very diverse genus, consisting of more than 350 species. As the genomic tools available to the shrub willow research community increase in both number and quality, so does the ability to characterize and deploy *Salix’s* innate diversity in improving high yielding cultivars. That diversity extends to variation between species even in traits as evolutionarily important as sex determination (Yang et al. 2021). By generating mapping populations that include characterized species crossed with those less studied, we will increase the number of *Salix* species available for trait introgression. We developed the *Salix* F_1_ HCP as a resource to characterize the interactions between alleles from different, but related species, and to map variation in important traits. Using GBS, we analyzed the population structure among the eight families, generated linkage maps of each of the parents using backcross markers and mapped the SDR in seven of the eight families using phenotypes collected from repeated field surveys. PCA and hierarchical clustering predominately resolved the population by section with the F_1_ clustering between the parents as expected. In each analysis, *S. alberti* P294 was found to very closely related to the *S. suchowensis* parents. Given these results and the limited publicly available information about *S. alberti*, P294 is likely *S. suchowensis* and will be described as such.

Among the 16 linkage maps produced, all QTL for sex were detected on chr15 in seven of the eight families and only in the maternal maps. In a recent study mapping sex in *S. triandra* using backcross markers, Li et al. (2020) were also only able to detect QTL within the maternal map, which is indicative of a ZW sex determination system on chr15. While this had been known for *S. purpurea* (Zhou et al. 2018), *S. viminalis* (Pucholt et al. 2017b), and *S. suchowensis* (Chen et al. 2016), this is the first study to report that both *S. koriyanagi* and *S. udensis* have ZW SDR on chr15. Although the location of the SDR has also not been reported in *S. integra*, the *S. integra* × *S. purpurea* family was entirely female and therefore was excluded in the linkage analysis. The chr15 map from *S. integra* P336 is considerably smaller than the other families, aligning to only the 14.2 – 15.5 Mb region of the *S. purpurea* reference genome, while maps of from of the other families had near complete coverage. The region *S. integra* P336 chr15 that did have segregating markers was outside the SDR intervals of the other female maps and is likely in the pseudoautosomal region of the sex chromosome.

Six of the eight families were female biased - a feature prevalent in *Salix*. Of those studied here, *S. purpurea* (Gouker et al. 2020), *S. viminalis* (Alström-Rapaport et al. 1997), *S. suchowensis* (Yang et al. 2020a), *S. udensis* (Ueno and Seiwa 2003), and *S. integra* (Tozawa et al. 2009) have documented cases of sex ratio bias, yet this is the first time it has been reported in *S. koriyanagi*. The genetic basis of sex ratio bias could be a result of secondary sex dimorphisms, such as higher mortality rates in males, increased herbivory and pathogen resistance in females, or the presence of a sex distorter locus (Pucholt et al. 2017a).

The mapping of the SDR in this study resulted in marker associations that represented a majority of chr15 prior to refinement. Of the four families with *S. purpurea* as the common female parent, refined QTL shared similar ranges of the SDR described by Zhou et al. (2020) (6.8 Mb in length starting at 2.3 Mb) with the exception of the *S. purpurea* × *S. viminalis* family (1.17 Mb). Carlson et al. (2019) mapped the SDR to 4.5 – 11.4 Mb on chr15 using a *S. purpurea* F_2_ population. This study localized the SDR to the same pericentromeric region of chr15 in *S. purpurea* using F_1_ hybrid families.

The most recent delimitation of the SDR in *S. viminalis* spanned roughly 3.4 Mb (approx. 2.3 – 5.7 Mb) of chr15 (Almeida et al. 2020). Our mapping of the SDR overlaps this region by 2.75 Mb, including the position of our peak marker, even though we aligned markers to a different reference genome. Almeida et al. (2020) aligned the SDR map to chr15 of both *S. purpurea* v.1.0 and *S. viminalis* reference genomes and found overall synteny between species, yet with several structural rearrangements. This contrasts with what is seen in our results, where the chr15 of 07-MBG-5027 showed only minor rearrangements, likely due to alignment to the high-quality *S. purpurea* v5.1 reference genome.

Based on annotations from the *Populus trichocarpa* genome, differential gene expression analysis in *S. suchowensis* between male and female plants led to predictions that the SDR was originally on chr14 (Liu et al. 2013). However, later work repositioned the SDR to the centromeric region of chr15 when based on *Salix* alignment (Chen et al. 2016). Our study defined the physical distance of the SDR at 2.44 – 9.35 Mb in P295 and 8.63 – 10.22 Mb in P294 with an overlap of 0.72 Mb. These two linkage maps show the greatest amount of rearrangement on chr15 when aligned to the *S. purpurea* reference. It is fair to conclude that alignment to a future *S. suchowensis* reference genome would aid in improving the mapping resolution in this region.

Suppressed recombination is a hallmark of chromosomes containing an SDR. Comparing the map of chr15 from each family, the SDR extends across a region of sparce marker density, approximately 3 to 9 Mb and is flanked by regions with greater marker density. As described above, this centromeric region with suppressed recombination is often associated with the SDR in *Salix*. In all seven families with mapped SDR, the peak marker was located within this region although its position varied. In the two *S. viminalis* families and two of the three *S. suchowensis* families, the peak marker was located within a 0.38 Mb region (2.57 – 2.95 Mb), while the remaining three families were less consistent and located proximal to the centromere. The generation of additional reference genomes for use in mapping will add context to these results and further refine the structure of the SDR among variable species.

This study described the population structure among the eight families within the *Salix* F_1_ HCP, constructed linkage maps for each parent, and mapped the SDR to the maternal chr15 in seven of the eight families. The introduction of the *Salix* F_1_ HCP provides the opportunity to map QTL for phenotypic traits beyond sex determination, while the linkage maps could be used to anchor and scaffold contigs in the generation of new reference genomes for each of the parents. While all species have a ZW sex determination system with an SDR that maps to chr15, these genetic resources provide a foundation for further characterization of the mechanism of sex determination and mapping of other key traits in these related species.

## Data Availability

The GBS data used in the population structure analysis and the eight files used for linkage map construction each family prior to linkage map construction are available in hapmap format through figshare at https://figshare.com at “GBS Data from Wilkerson et al”. R code used to format and create the linkage maps is available on the Willowpedia Github site located at https://github.com/Willowpedia/Wilkerson_etal_SalixLinkageMaps. Table S1 contains statistics on each of the 16 linkage maps created in this study, including marker count and cM length for each linkage group and the total markers and cM length for each map. Figure S1 is a pdf slide show that, one map per slide, shows the marker cM by physical position for each linkage group and a heatmap visualizing recombination frequency and linkage. Tables S2 and S3 contain the cM distance and sex QTL LOD scores of markers within each female and male map, respectively.

## Acknowledgements

We are grateful for the excellent technical support contributed by Lauren Carlson, Jane Petzoldt, Dawn Fishback, and Rebecca Wilk.

